# Experimental Parameters-Based Monte-Carlo Simulation of Single-Molecule Localization Microscopy of Nuclear Pore Complex to Evaluate Clustering Algorithms

**DOI:** 10.1101/2022.09.21.508613

**Authors:** Wei-Hong Yeo, Yang Zhang, Amy E. Neely, Xiaomin Bao, Cheng Sun, Hao F. Zhang

## Abstract

Single-molecule localization microscopy (SMLM) enables the detailed visualization of nuclear pore complexes (NPC) *in vitro* with sub-20 nm resolution. However, it is challenging to translate the localized coordinates in SMLM images to NPC functions because different algorithms to cluster localizations as individual NPCs can be biased without ground truth for validation. We developed a Monte-Carlo simulation to generate synthetic SMLM images of NPC and used the simulated NPC images as the ground truth to evaluate the performance of six clustering algorithms. We identified HDBSCAN as the optimal clustering algorithm for NPC counting and sizing. Furthermore, we compared the clustering results between the experimental and synthetic data for NUP133, a subunit in the NPC, and found them to be in good agreement.

## Introduction

The nuclear pore complex (NPC) forms a nanosized channel on the nuclear membrane. It is an essential gateway for transporting biomolecules (e.g., RNA, proteins) between the cell nucleus and the cytoplasm [1,2]. The NPC has highly preserved ring-shaped features across different mammalian species and is composed of about 30 different nucleoporin subunits (NUPs) in human cells with well-defined NUP stoichiometries [1]. The scaffold NUPs, are typically classified into outer-ring, inner-ring, and transmembrane NUPs (Figure 1) and have either 8 or 16 copies of the corresponding NUPs possessing 8-way symmetry. While state-of-the-art structural biological techniques (e.g., X-Ray crystallography and cryogenic electron microscopy) provide the detailed structural organization of NPCs [3], the functional information of specific NUPs under different cellular conditions remains understudied.

**Figure 1.**
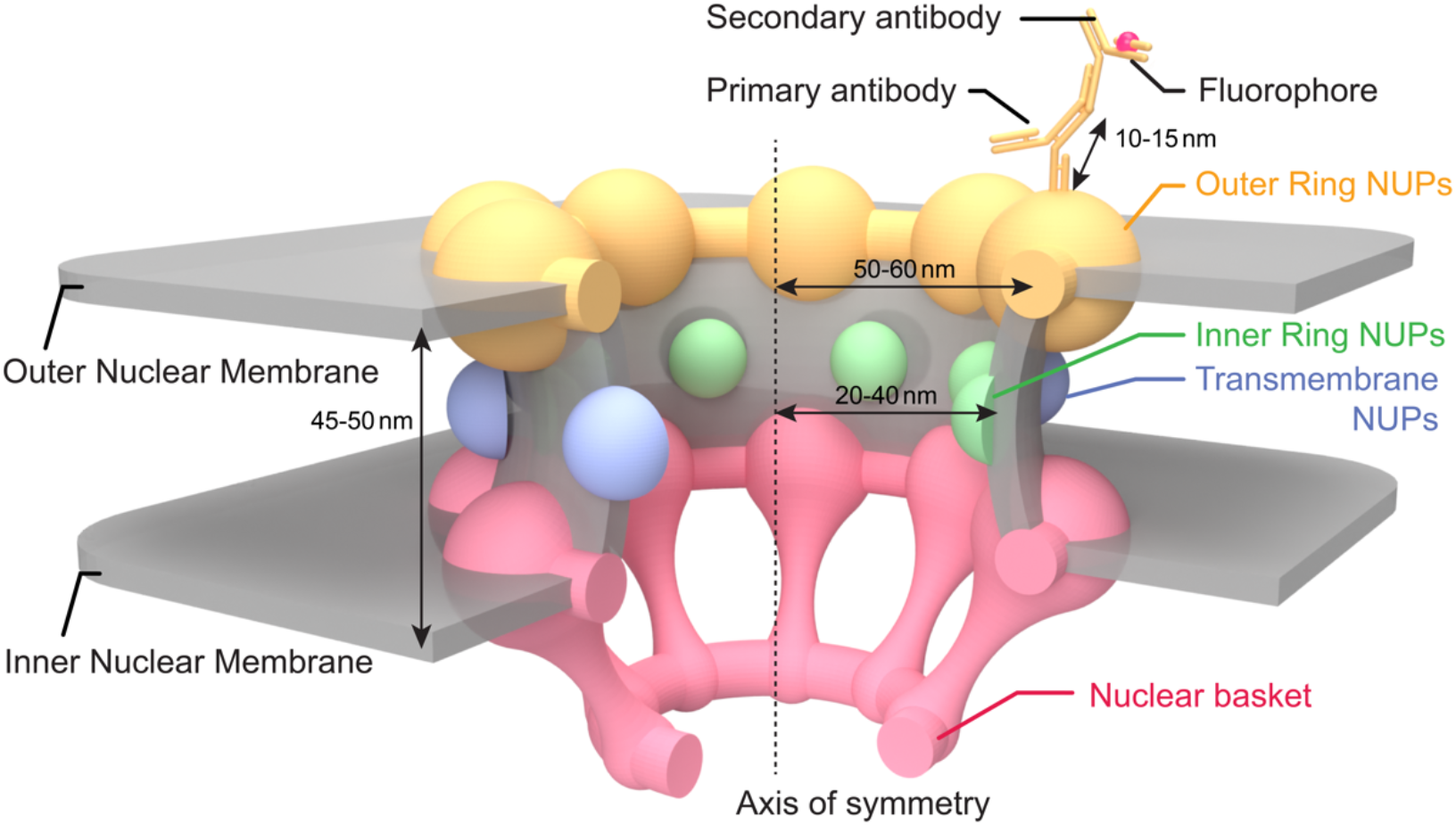
Illustration of the physical sizes and relationship between the structural components of NPC, outer ring NUPs, inner ring NUPs, transmembrane NUPs, and nuclear NUPs. On the outer ring NUP, we show fluorescence labeling by a primary antibody and a secondary antibody with a fluorophore.

Single-molecule localization microscopy (SMLM) allows optical imaging of subcellular architectures at the molecular level based on stochastic fluorescence emissions with high molecular specificity [4,5]. SMLM enables the capture of the otherwise inaccessible spatial and functional information of biomolecules [6–9]. In NPC imaging, NUPs have not only been visualized using SMLM [10] but have also been treated as reference standards for benchmarking the imaging performance of various SMLM techniques [11] because of NPC’s well-defined and highly-preserved nanostructures. However, the objective analysis of functional information in NPC presented in SMLM images remains challenging. This is because it requires clustering of the coordinates data obtained from SMLM into individual NPCs before investigating any NPC functions (e.g., NUP composition, NPC size & density). Existing clustering methods typically have a biased clustering outcome due to the subjective selection of clustering parameters (e.g., minimum size and points of the cluster). Using ground-truth information would allow us to optimize and validate the clustering performance [12]; however, such ground-truth data is not readily available in SMLM imaging of NPC.

To overcome this challenge, we developed a Monte-Carlo simulation method to simulate NPC images following the physical process in SMLM experiments to generate ground-truth NPC data. The simulation incorporates several fundamental physical uncertainties that collectively determine each single-molecule localization (SML) coordinate, thereby providing well-controlled and realistic subjects for NPC clustering. First, in the SMLM data, each NPC will be visualized as a point cloud consisting of the coordinates of SMLs. The SMLs arise from the stochastic fluorescence photoswitching events of fluorescent labels (marked as the small red symbol on the secondary antibody in Figure 1) that are tagged to selected NUPs [4]. Second, we considered the influence of uncertainties arising from indirect immunofluorescence labeling [8]. Because the antibodies used to label NPCs often have an inherently large size (∼10-15 nm), localizing the fluorescence molecules entangles complicates the true localization uncertainty of target molecules in SMLM [8]. Second, the flexible Y-shaped structure of antibodies makes it even more challenging to correlate the localization events with the true location of the target molecule as antibodies may orient in different directions. Third, we introduced localization uncertainties and realistic non-specific SML background from antibody off-targeting and background autofluorescence signals [13] in the final SMLM image. By providing a model that links the ground-truth geometry to an SMLM image, this work sets the stage for SMLM to validate and optimize clustering methods for objective NPC clustering and subsequent functional analyses.

## Results and discussion

### SMLM imaging of NUP133 using indirect immunofluorescence labeling

We first statistically characterized three experimental parameters of NPCs to be used in our Monte-Carlo simulation: (1) localization precision, (2) SMLs per labeling site, and (3) background SMLs. We imaged NUP133, an outer-ring NUP with an average radius of 53.5 nm to the central axis, using our home-built experimental SMLM system [14]. The SMLM image of NUP133 shows individual clusters spanning the entire nuclear membrane (Figure 2A). The magnified view shows the characteristic nanosized ring-shaped features (Figure 2B). Particularly, only a portion of the 8 NUP133 proteins was labeled and visualized in each NPC the steric hindrance of the antibodies mask the reactive sites for the antigen-antibody reactions, and intrinsically poor labeling efficiency of the primary and secondary antibodies. We observed smaller localization clusters (highlighted by the arrows in Figure 2B) which could be detected as the SMLs from (1) true individual NPCs signals at relatively low labeling efficiency (*i*.*e*., 1 of 8 NPC sites labeled) or (2) non-specific binding of secondary antibodies. We obtained an average localization precision of 15 nm measured from ∼70,000 SMLs, and the localization precision follows a normal distribution (Figure 2C). We further deposited the dye-labeled secondary antibody on a glass substrate and found that the SMLs per antibody follows an exponential distribution with a mean value of 10 (Figure 2D), which was used to simulate the number of SMLs from each NUP protein. Our imaging results of the NUP133 protein are consistent with the literature [10].

**Figure 2.**
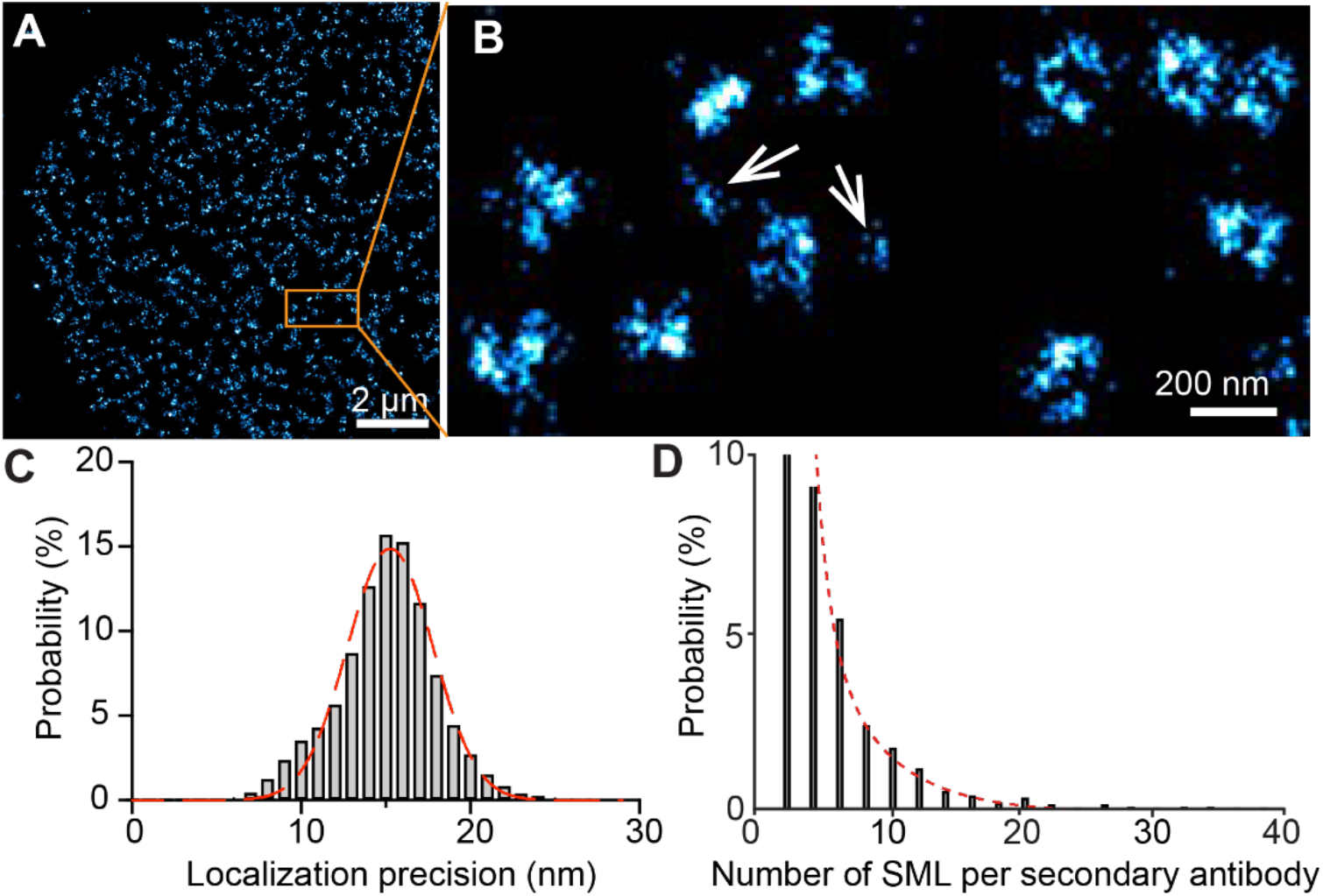
(A) SMLM imaging reconstruction of NUP133 labeled HeLa cells; (B) a magnified view of the area highlighted by the orange box in panel A; (C) A histogram showing the probability distribution of localization precision from ∼70,000 single molecules detected in panel A and a fitted curve using a gaussian distribution (red curve); (D) A histogram showing the probability distribution of the number of SMLs in Alexa Fluor 647-labeled secondary antibody fitted onto an exponential distribution (red curve).

The lack of ground truth information makes it challenging to identify true NPC clusters visually, as shown in the cases in Figure 2B. Therefore, our goal is to comprehend the major sources of uncertainty to simulate the NPC images resembling the experimental conditions. We intentionally used the classical indirect immunofluorescence labeling method (i.e., an anti-NUP133 primary antibody and the corresponding Alexa Fluor 647-tagged secondary antibody) to label the NUP133 protein to present a challenging case with multiple sources of uncertainties. Thereafter, we incorporated these variations into our Monte-Carlo simulation in addition to the localization uncertainties.

### Monte-Carlo simulation of SMLM imaging of NPC

Using the experimentally acquired probability density functions (PDFs) of localization precision and number of SMLs from each secondary antibody (Figures 2C-2D) and the physical size of primary and secondary antibodies, we summarized the overall Monte-Carlo simulation process with the four main functions described in the flowchart (Figure 3). First, we simulated the spatial distribution of NPCs and the coordinates of NUPs of interest. NPCs are known to be randomly distributed on the nuclear membrane with varying pore distances of approximately 250 to 500 nm [15]. We simplified the spread model of NPC as projections on a two-dimensional (2D) squared box with varying NPC spatial density based on the following two rationales: (1) most SMLM studies of NPC use TIRF illumination to selectively image the NPCs close to the flat cell-glass interface, and (2) the typical axial resolution of ∼80 nm in SMLM is challenging to resolve the NUPs on the nuclear membrane. Then, we obtained the distance-to-center parameter of different NUPs from the literature [3]. We assumed that all the NUPs maintained 8-way symmetricity and generated eight sites around the NPC centroid (orange dots in Figure 4A). Next, we used the Poisson disk algorithm [16] to generate the NPC centroid locations (green circles with black crosses in Figure 4A) with a minimum spacing of 250 nm and a distance of 53 nm to simulate NUP133 as an example. We then trimmed the list of candidate NPC centroids to 1-9 NPCs per μm^2^, corresponding to the NPC densities reported in typical mammalian cells [17].

**Figure 3.**
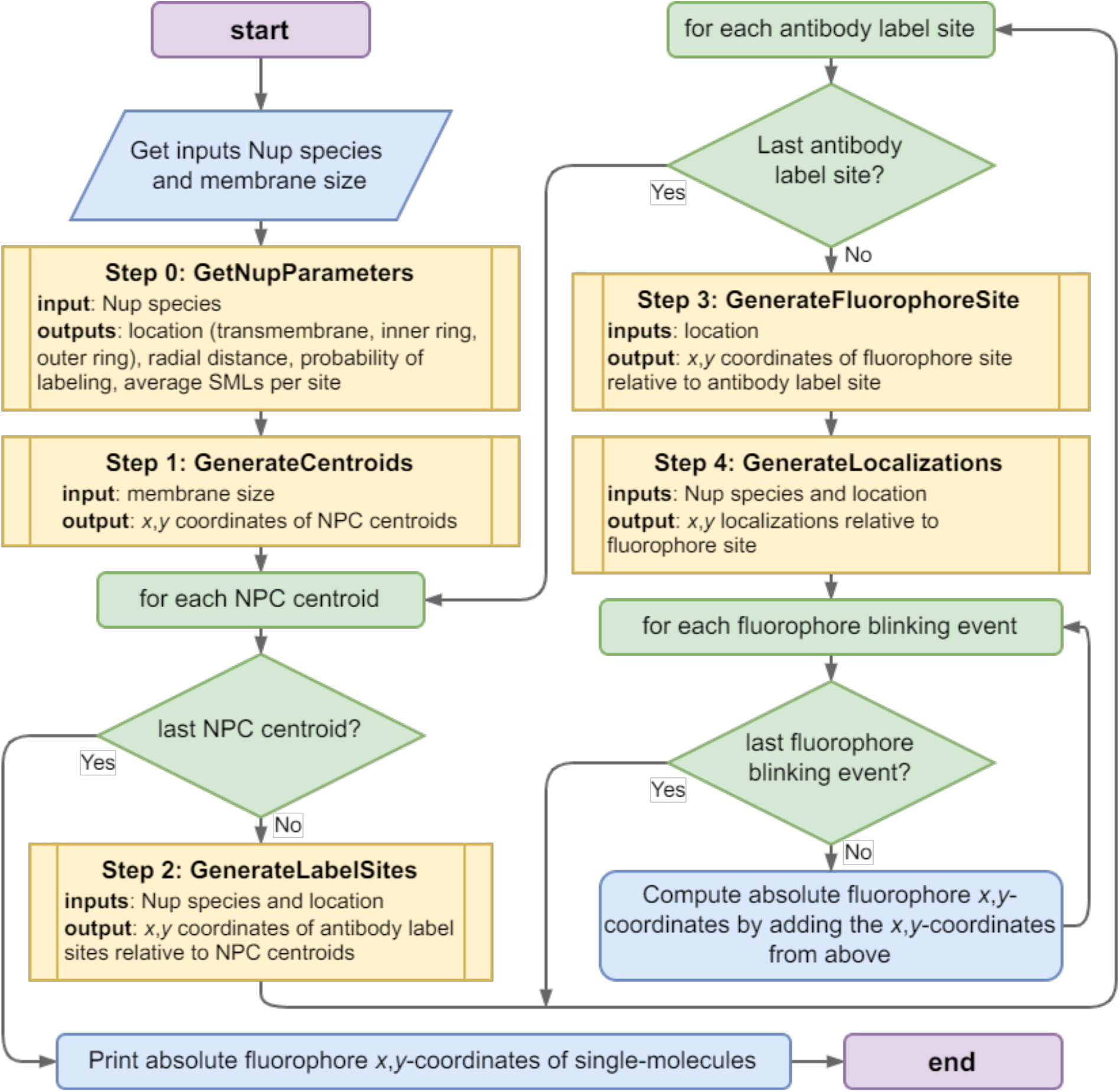
Flowchart of our Monte-Carlo simulation. GetNupParameters, GenerateCentroids, GenerateLabelSites, GenerateFluorophoreSite, and GenerateLocalizations are functions used in the simulation and can be found at https://github.com/FOIL-NU/MC-NPC.

**Figure 4.**
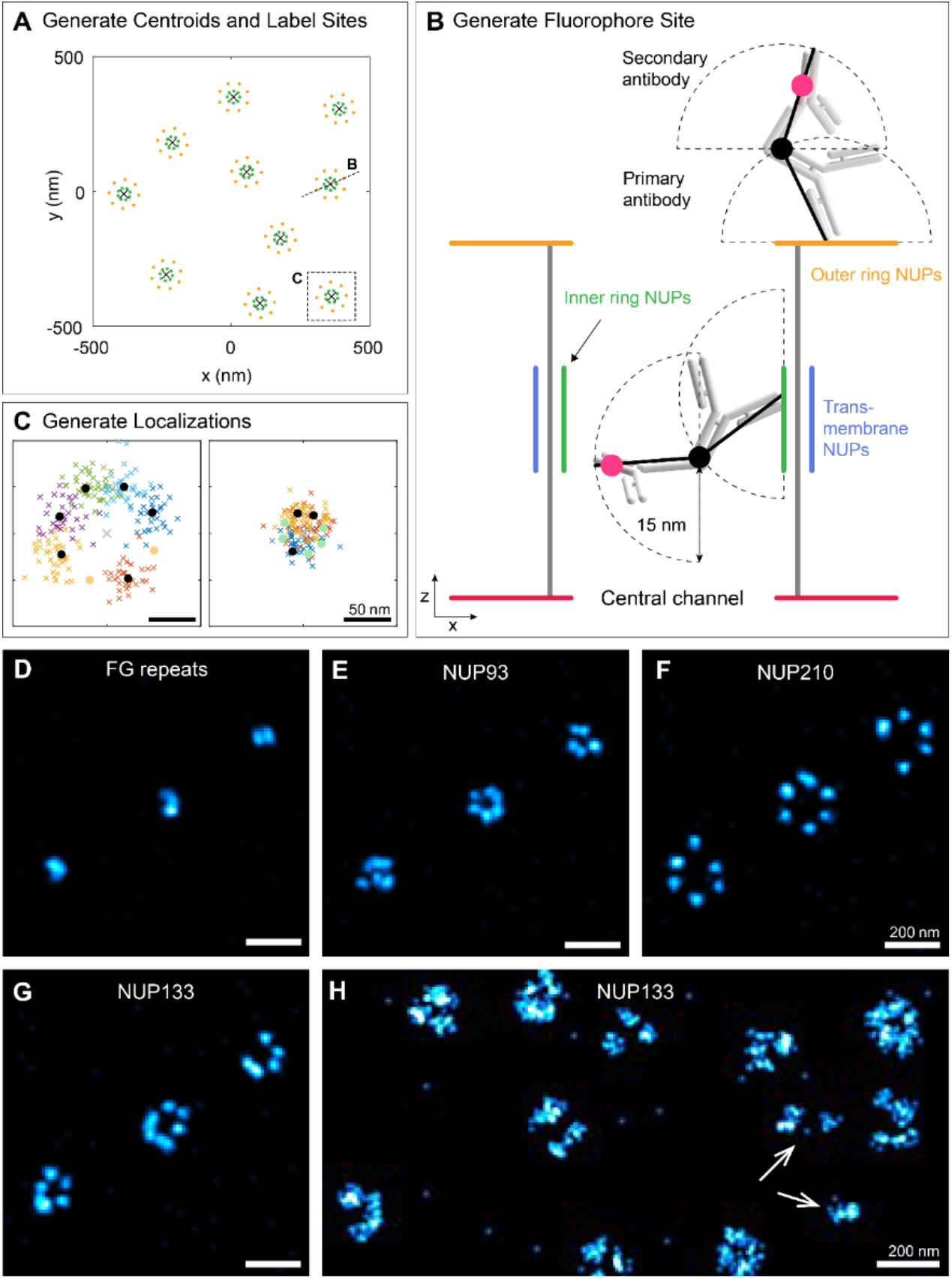
(A) An illustration of the initial centroid and label site generation process. Black crosses: centroids generated; yellow and green circles: label sites of outer ring NUPs and inner ring NUPs, respectively; (B) A illustrated cross-section of a single NPC site with dotted hemispherical regions depicting possible locations of antibodies due to the degree of freedom with labeling. Fluorophores are randomly assigned along the length of the secondary antibody (pink dots); (C) A detailed illustration of a single NPC site. Scatterplots of the coordinates simulating a single NPC for an outer NUP (left) and inner ring (right). Colored crosses: SMLs; black dots: primary antibody; green circles and gray crosses indicate the center location of the NPC as references; (D) Simulated SMLM images of FG repeats; (E) Simulated SMLM images of NUP93; (F) Simulated SMLM images of NUP210; (G) Simulated SMLM images of NUP133 with 3 NPCs/μm^2^; (H) Simulated SMLM images of NUP133 using the identical experimental uncertainties in Figure 2.

Second, we simulated the effects of indirect immunofluorescence labeling on different NUPs. On the one hand, antibodies are known to show finite labeling efficiency to the protein of interest because of antibody affinity [18] and steric hindrance around the proteins [19]. Such labeling inefficiencies have been observed in SMLM-NPC imaging, with <50% of the NUPs being labeled and visualized [20]. On the other hand, antibody labeling of NUPs can be anisotropic, such as the labeling of inner-ring NUPs and transmembrane NPCs. More specifically, the antibodies may only access the reactive sites of the inner-ring NUPs from the central channel region because the other side of the inner-ring NUP is blocked by the transmembrane NUPs (Figure 4B). The linker length, coupled with the labeling directionality can cause underestimates of the radius of the inner NUP rings. A similar effect has been observed in SMLM imaging of microtubules, wherein antibodies can only access the outer wall of the microtubule and cause an elevated diameter measurement [21]. In contrast, antibodies can presumably access the outer ring NUPs isotropically because they are labeled parallel to the central channel (Figure 4B). To simplify the model of the antibodies, we assume that primary and secondary antibodies are 15 nm rigid structures that form two spheroid joints on the target proteins (Figure 4B). We assumed that they can fall within two hemispheres orthogonal to the target site since it is likely difficult for the secondary antibody to fold back onto the primary antibody due to steric hindrance. We also assumed that secondary antibodies only bind to the primary antibody’s farthest end, representing the worst-case scenario of labeling for SMLM. Further, we assumed that fluorophores are uniformly distributed along the length of the secondary antibodies. Most fluorophores are bioconjugated to the antibodies through covalent reactions to the largely abundant amino, carboxyl, or thiol residues on the proteins, which possess similar reactivities [22]. Collectively, we incorporated antibody accessibility and used a binomial distribution on the eight sites for labeling efficiency to generate the labeling sites of the NUPs.

Third, for each fluorophore site generated, we used a random number of SMLs according to the experimentally measured PDF of SMLs. We assumed a uniform localization uncertainty of 10 nm for SMLs to represent the 20 nm resolution [23] unless otherwise stated. The overall simulation outcome for each NPC is a point cloud representing the SMLs typically detected in the SMLM images (Figure 4C). A representative NUP133 simulation shows the labeling of six sites of the eight subunits, which can readily be resolved visually (left panel in Figure 4C). In contrast, as the anisotropic labeling of an inner-ring NUP, the three labeled sites are unresolved after convolving with the experimental PDFs (right panel in Figure 4C). With the simulation, we can trace the origins of each SML from the individual NUPs (shown as the six different colored point clouds in Figure 4C), which would be challenging to identify without knowing the ground truth.

Using the coordinate list output from the Monte-Carlo simulation, we rendered representative synthetic SMLM images, FG repeats (central channel NUPs), NUP93 (inner-ring NUPs), NUP210 (transmembrane NUPs), and NUP133 (outer-ring NUPs) (Figures 3D-3G). These four NUPs have a radius of 53 nm, 10 nm, 40 nm, and 80 nm, respectively. NUP210 can be fully resolved using antibodies because of the relatively large separation among individual NUP210 proteins along with its labeling direction (Figure 4F), which is consistent with the published experimental results [24]. However, the visualization of the ring-shaped structures for NUP93 and FG-repeats is ambiguous (Figures 3D-3E), suggesting that only the transmembrane or outer-ring NUPs are resolvable in SMLM using an indirect immunofluorescence labeling approach. We also simulated the NUP133 SMLM image (Figure 4H) with the experimental PDFs. The overall quality of the image visually resembles the experimental results. In both images, ten NPCs are unambiguously identified with diameters of ∼100 nm with their unique ring-shaped structures. In addition, two smaller clusters (highlighted by the arrows in Figure 4H) arose from extremely low-efficiency labeling of an NPC. In short, these SMLM images generated by Monte-Carlo simulation can reflect the structural organization of different NUPs and are consistent with our experimental results.

### Clustering performance test using simulated NPCs

Treating the Monte-Carlo simulated NPCs as the ground truth, we investigated the optimal clustering method for NPC for SMLM. We selected density-based spatial clustering of applications with noise (DBSCAN) [25], Hierarchical DBSCAN (HDBSCAN) [26], Ordering points to identify the clustering structure (OPTICS) [27], Agglomerative [28], K-means [29], and balanced iterative reducing and clustering using hierarchies (BIRCH) [30] methods in our test. While the first three do not require the input of the number of clusters as a parameter, the latter three require it. To systematically test their clustering performance, we generated synthetic SMLM images of NUP133 with a relatively high NPC density of 9 NPC per µm^2^, which is among typical values reported for mammalian cells [17] to present challenging cases. We generated a 10×10 µm^2^ area of NPCs with NUP133 as labeling targets (Figure 5A). As we assume that our labeling efficiency follows a binomial labeling process, NUP133 clusters have diverse shapes with 2-6 sites labeled (magnified views in Figure 5A).

**Figure 5.**
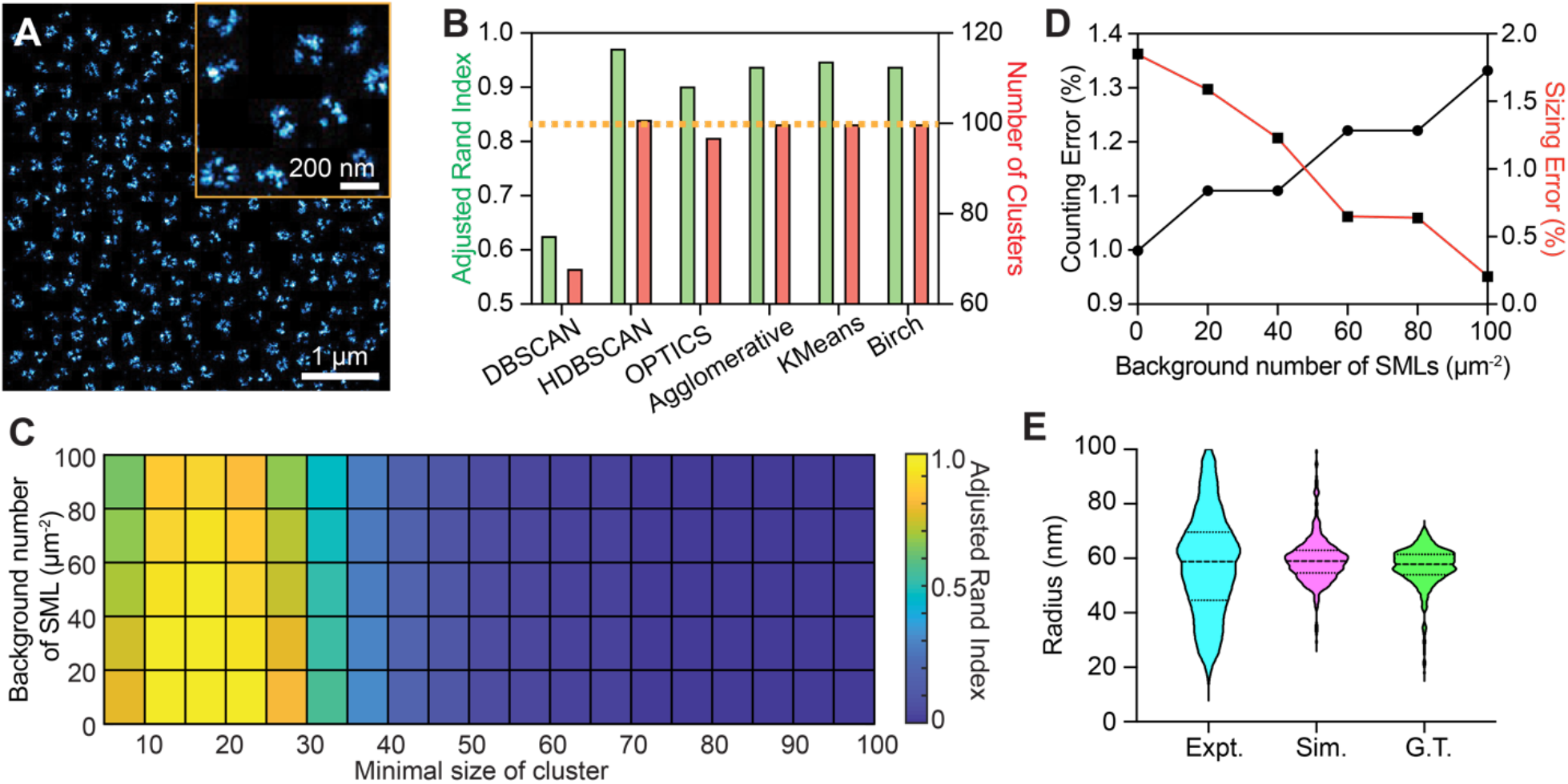
(A) A simulated SMLM image of NUP133 with 9 NPCs/μm^2^ with background levels of 100 SMLs per μm^2^; (B) Adjusted Rand Index (green bars) and the number of clusters (red bars) values using different clustering algorithms. The dashed line indicates the ground-truth number of clusters; (C) A heatmap showing the Adjusted Rand Index for clustering performance characterization as a function of increased background number of SML and minimal size of the cluster in the HDBSCAN parameter; (D) The counting (black line) and sizing (red line) errors of HDBSCAN clustering at a background number of 40 SMLs, with a minimum cluster size of 15 using the image shown in panel A; (E) A violin plot illustrating the histogram radius distribution of the NUP133-ring of the experimental data (cyan) and the simulated data (magenta) of each NUP133 cluster after HDBSCAN clustering and using the ground-truth (green).

We used an Adjusted Rand Index using Python’s scikit-learn package to quantitatively compare the clustering performance of the tested methodsThe Adjusted Rand Index calculates the similarity between clustering results and the ground truth, and is used extensively for characterizing clustering performance [12]. An Adjusted Rand Index approaching 1 indicates better clustering performance (Figure 5B), and with this, we found that HDBSCAN stands out among all the other algorithms tested (green bars in Figure 5B). Additionally, the number of clusters identified from the six algorithms are 55, 101, 90, 100, 100, and 100 for DBSCAN, HDBSCAN, OPTICS, Agglomerative, KMeans, and BIRCH, respectively (red bars in Figure 5B). Notably, as Agglomerative, KMeans, and Birch require the number of clusters in the dataset to be provided, we used a value of 100, which matched the ground truth (as indicated with the dashed yellow line in Figure 5B) to test their theoretical clustering performances. The NPC counting percent errors were then calculated with (N_d_−N_g_)/N_g_×100%, where N_d_ is the number of NPC detected by the algorithm, and N_g_ is the number of NPC in the ground truth, and the numbers are 45%, 1%, 10% for DBSCAN, HDBSCAN, and OPTICS, respectively. The percent errors of the other three algorithms were not calculated because we used N_d_ = N_g_ = 100, which give percent errors of 0%. However, it is practically challenging to select a number-of-clusters value that exactly matches the ground truth in these three algorithms. Furthermore, considering the high Adjusted Rand index available in HDBSCAN without the need to input the number-of-clusters value, we conclude that HDBSCAN is the optimal clustering algorithm for NPC identification.

Next, we investigated HDBSCAN’s sizing and counting performance under different background SMLs and NPC density conditions, which might affect the clustering performance. The number of SMLs per NPC and the background count of SMLs in the simulations follow experimental parameters (Table 1). We compared the Adjusted Rand Index values to test the clustering performance in this case. As the minimal cluster size is the only required input parameter in HDBSCAN, we first optimized this parameter in our clustering performance at different background SML levels. Concurrently varying the minimal cluster size with the background levels, we observed that a cluster size of 15 yielded the highest Adjusted Rand Index (∼0.9–1.0) at different background SML levels, as shown in Figure 5C as a heatmap. Additionally, the heatmap’s second to fourth columns in Figure 5C illustrates minimal variations of the Adjusted Rand Index at different background levels. This feature highlights the robustness of HDBSCAN in NPC clustering without prior knowledge of the number of clusters and with different background conditions that are potentially associated with experimental biological imaging conditions.

**Table 1.** Summary of NUP parameters used in the simulation.

| NPC protein | Location | Radius relative to NPC centroid (nm) |
| --- | --- | --- |
| NUP93 | Inner Ring | 40.0 |
| NUP133 | Outer Ring | 53.5 |
| NUP210/GP210 | Transmembrane NUP | 80.0 |
| FG repeats | Central Channel | 10.0 |

Further, we calculated the counting and sizing errors to better understand the clustering performance (Figure 5D). Specifically, the counting errors increase from 1.0% to 1.3%, while the sizing errors decrease from 1.9% to 0.3% as a function of increasing background SMLs. Consequently, these errors are relatively small in understanding the potential change in NPC functions of size and density variations. The minimal variations in sizing and counting errors at different background levels suggest that HDBSCAN is unlikely to be biased when analyzing different biological samples with various autofluorescence and non-specific binding conditions. Finally, we quantified the NUP133 cluster size using the experimental, simulated, and ground truth datasets extracted from HDBSCAN (Figure 5E). The average radius of the NUP133 clusters are 58 nm, 59 nm, and 59 nm for the experimental, simulated, and ground-truth datasets, respectively. These values are consistent with the reported 53 nm radius plus the antibody labeling and localization uncertainties [11].

## Methods

### 1. Sample preparation and SMLM imaging

HeLa cells (ATCC) were grown in DMEM (Gibco/Life Technologies) supplemented with 2 mM L-glutamine (Gibco), 10% fetal bovine serum (Gibco/Life Technologies), and 1% penicillin / streptomycin (10 kU/mL, Gibco/LifeTechnologies) at 37°C with 5% CO_2_. The cells were plated on a No. 1 borosilicate bottom 8-well Lab-Tek™ Chambered Cover Glass with 30%-50% confluency. After 48 hours, the cells were permeabilized and fixed following the literature protocol [10]. The fixed samples were blocked with a blocking buffer (3% BSA, 0.5% Triton X-100 in PBS) for 20 min and then incubated with the primary antibodies (Rabbit anti-NUP133, 1:100 dilution, Sigma HPA059767) in the blocking buffer overnight at 4°C and rinsed with a washing buffer (0.2% BSA, 0.1% Triton X-100 in PBS) thrice. The samples were further incubated with the corresponding Donkey secondary antibodies-dye conjugates (Anti-Mouse Alexa Fluor 647, 2.5 μg/mL in blocking buffer) for 40 min at 25°C, washed thoroughly with PBS thrice at 25°C, and then stored at 4°C. SMLM imaging was performed with a custom-built SMLM system based on a Nikon Ti-2E inverted fluorescence microscope, as described before [31]. The samples were immersed in an SMLM imaging buffer [32], excited with a 647 nm continuous-wave laser under total internal reflection fluorescence (TIRF) illumination mode. An exposure time of 20 ms was used and an image sequence of 20,000 frames was collected to reconstruct the SMLM image using the ThunderSTORM plugin in ImageJ [33].

### 2. Monte-Carlo simulation

The Monte-Carlo simulation [34] was implemented in MATLAB 2022a with main functions following four main steps shown in Figure 3. In the first step, we used the Poisson disk algorithm [16] to generate the NPC centroid locations with a minimum spacing of 250 nm. The number of centroids was then adjusted to give the desired NPC density. With each NPC centroid location, the label sites were simulated using a uniform distribution between [0°, 45°) to simulate the rotational freedom of the nuclear pore complex within the nuclear membrane.

In the second step, we assigned a probability on each of the eight rotationally symmetric NUPs sites depending on the NUP site (Table 2) to simulate the stochastic process of antibody binding onto each available NUP site.

**Table 2.** Summary of simulation constants unless otherwise noted.

| Parameter | Value |
| --- | --- |
| Localization uncertainty | 10 nm |
| Primary antibody length | 15 nm |
| Secondary antibody length | 15 nm |
| Chemical tag length | 5 nm |
| Probability of labeling onto a NUP133 protein | 0.3 per site |
| Average number of SMLs per secondary antibody | 10 |
| Nuclear membrane size | $10 \times 10 \mu\text{m}^2$ |
| Minimum spacing between adjacent NPCs | 250 nm |
| Background noise | 0, 40, and 100 photons per $\mu\text{m}^2$ |
| Density of NPCs | 1, 3, 5, 7, and 9 NPCs per $\mu\text{m}^2$ |

In the third step, we simulated the location of individual fluorophores on the antibody. To simplify the model of the antibodies, we assumed that primary and secondary antibodies were 15 nm rigid structures that form two spheroid joints on the target proteins (Figure 4B). For each antibody, we assumed that they fall within a hemisphere normal to the target site. The angle of attachment of each antibody was simulated using a uniform distribution for the azimuth with [0°, 360°) and a uniform distribution scaled by an inverse sine function between [0,1], giving an elevation of [0°, 90°]. This ensures that the points generated were equally dense throughout the semicircular region. We assumed that secondary antibodies only bind to the primary antibody’s farthest end, representing the worst-case scenario of labeling in imaging. Further, we used a uniform distribution between [0, 15] nm to generate the fluorophore location along the length of the secondary antibodies.

In the last step, we used a random number generator that follows a negative binomial distribution [35] to generate the number of SMLs with parameters adjusted so that the mean number matched our experimental results. For each SML, we assumed a uniform localization uncertainty of 10 nm, and we modeled the spread of the localizations using a symmetric bivariate normal distribution. The MATLAB code for our simulation is available on Github at https://github.com/FOIL-NU/MC-NPC.

### 3. Clustering test

We carried out all the clustering tests using the Python packages Scikit-learn 1.1.2 for DBSCAN, OPTICS, Agglomerative, KMeans, and BIRCH; and an individual package of HDBSCAN. When investigating the performance among different clustering methods, we selected consistent parameters, including a minimum point of 10 and a minimum size of 30, to minimize the chance of extracting repeatedly emitting individual fluorophores. In the cases of Agglomerative, KMeans, and BIRCH, we used the number of clusters of 100, which was identical to that of the ground truth, to test their theoretical performance. In the case of HDBSCAN, we only selected a minimum size of 30 as the rest of the parameters were automatically optimized by the intrinsic hierarchical clustering feature.

## Conclusion

In this work, we developed a Monte-Carlo model based on experimental SMLM parameters, structural organization of NPC subunits, uncertainties due to the physical sizes of molecular labelings, and the single molecule localization uncertainties due to stochastic blinking events. We used this model to generate ground truth NPC images and tested six reported clustering methods. We found that HDBSCAN was the optimal clustering algorithm for NPC, with the lowest errors. In addition, HDBSCAN provides a consistent measurement of the radius of NUP133 rings from both experimental and Monte-Carlo simulated images compared to the ground truth.

## Acknowledgments

The authors acknowledge the generous support from the NIH grants R21GM141675, R01EY026078, R01EY019949, R01GM140478, R01GM139151, R01GM143397, R01AR075015, and U54CA268084 and NSF grant CBET-1706642, CHE-1954430, and EFRI-1830969, and the American Cancer Society Research Scholar Grant RSG-21-018-01-DDC. Wei-Hong Yeo is supported by the Christine Enroth-Cugell Fellowship for Vison and Neuroscience at Northwestern University.

## Notes

https://github.com/FOIL-NU/MC-NPC

### Competing Interest Statement

The authors have declared no competing interest.

